# An evolution-based high-fidelity method of epistasis measurement: theory and application to influenza

**DOI:** 10.1101/2019.12.11.873307

**Authors:** Gabriele Pedruzzi, Igor M. Rouzine

**Affiliations:** Sorbonne Université, Institute de Biologie Paris-Seine, Laboratoire de Biologie Computationelle et Quantitative LCQB, F-75004, Paris, France

**Author notes:** Corresponding author: Igor Rouzine, Evolution and immunology of pathogens, Laboratory of Computational and Quantitative Biology, 7238 CNRS-UPMC, Bâtiment C, C311C, Sorbonne Université Campus Pierre et Marie Curie, 4 rue Jussieu, Paris 75005, http://www.ibps.upmc.fr/fr/Recherche/umr-7238/evolution-immunologie-pathogenes. Influenza protein sequence data are from https://www.fludb.org.

**Keywords:** mathematical model, quasi-equilibrium, false-positive, linkage

## Abstract

Linkage effects in a multi-locus population strongly influence its evolution. The models based on the traveling wave approach enable us to predict the speed of evolution and the statistics of phylogeny. However, predicting the evolution of specific sites and pairs of sites in the multi-locus context remains a mathematical challenge. In particular, the effects of epistasis, the interaction of gene regions contributing to phenotype, is difficult both to predict theoretically and detect experimentally in sequence data. A large number of false interactions arise from stochastic linkage effects and indirect interactions, which mask true interactions. Here we develop a method to filter out false-positive interactions. We start by demonstrating that the averaging of the two-way haplotype frequencies over a multiple independent populations is necessary but not sufficient, because it still leaves high numbers of false interactions. To compensate for this residual stochastic noise, we develop a triple-way haplotype method isolating true interactions. The fidelity of the method is confirmed using simulated genetic sequences evolved with a known epistatic network. The method is then applied to a large database sequences of neurominidase protein of influenza A H1N1 obtained from various geographic locations to infer the epistatic network responsible for the difference between the pre-pandemic virus and the pandemic strain of 2009. These results present a simple and reliable technique to measure site-site interactions from sequence data.

**Author’s summary:** Interaction of genomic sites creating “fitness landscape” is very important for predicting the escape of viruses from drugs and immune response and for passing through fitness valleys. Many efforts have been invested into measuring these interactions from DNA sequence sets. Unfortunately, reproducibility of the results remains low, due partly to a very small fraction of interaction pairs, and partly to stochastic noise intrinsic for evolution masking true interactions. Here we propose a method based on analysis of genetic sequences at three genomic sites to clean stochastic linkage and apply it to influenza virus sequence data.

## Introduction

Almost a century ago, it was realized that the evolution of a population is strongly affected by the fact that the fates of alleles at different loci are linked unless separated by recombination. These *linkage* effects include clonal interference (Fisher 1930; Muller 1932), background selection, genetic hitchhiking (Rice 2004), enhanced accumulation of deleterious mutations (Muller ratchet) (Felsenstein 1974), and the increase of genetic drift on one locus due to selection at another (Hill and Robertson 1966). Linkage decreases the speed of adaptation and creates random associations between pairs of mutations occurring on the same branch of the ancestral tree.

These effects have been taken into account in early mathematical models considering two loci (KIMURA 1994) and, more recently, in the traveling wave approach, which describes an arbitrarily large number of connected sites (Rouzine *et al*. 2003; Desai and Fisher 2007; BRUNET *et al*. 2008; Rouzine *et al*. 2008; Hallatschek 2011; Good *et al*. 2012). These models describe the dynamics of fitness classes and include the factors of selection, mutation, random genetic drift and, sometimes, recombination (Rouzine and Coffin 2005; Neher *et al*. 2010; Rouzine and Coffin 2010). All these models predict a narrow fitness distribution traveling in the fitness space in a direction depending on the initial conditions and parameters. This “traveling wave” consists from the deterministic bulk and the leading stochastic edge, where generation and establishment of rare beneficial mutations limit the adaptation rate. Alternatively, the distribution may move backwards accumulating more and more deleterious alleles (Muller ratchet). These models are able to express, in the general form, important observable quantities in terms of model parameters, such as the population size, mutation rate, and the distribution of selection coefficients over loci. The observable quantities include the adaptation rate (Rouzine *et al*. 2003; Desai and Fisher 2007; Brunet *et al*. 2008; Rouzine *et al*. 2008), Muller ratchet rate (ROUZINE *et al*. 2003; Rouzine *et al*. 2008), the conditions of full equilibrium (Rouzine *et al*. 2003; Goyal *et al*. 2012), fixation probability of an allele, and the most probable selection coefficient (Good *et al*. 2012). The same general approach was used to predict the statistical properties of the ancestral tree (Brunet *et al*. 2007; Rouzine and Coffin 2007; Rouzine and Coffin 2010; Walczak *et al*. 2012; Neher and Hallatschek 2013).

Despite of all the progress, prediction of the evolution of specific sites in the multi-site context remains an open question. How do allelic frequencies at each site change in time when the system is adapting? Although they are known to follow random trajectories, what can be said about the average allelic frequency of a given site with a given fitness effect of mutation? Also, what can we say about the evolution of site pairs, especially in the presence of epistatic interaction?

Epistasis, defined as co-selection, the interaction of genes and gene regions contributing to phenotype, is an omnipresent phenomenon (Carlborg *et al*. 2006). Gene interactions are reported to be responsible for a considerable fraction of the organism’s genetic inheritance (Zuk *et al*. 2012). They create fitness valleys that the evolutionary path has to cross (Weissman *et al*. 2009). In pathogens, epistasis facilitates the development of drug resistance and immune escape and impedes reversion of resistant mutations (Nijhuis *et al*. 1999; Levin *et al*. 2000; Piana *et al*. 2002; Handel *et al*. 2006; Gonzalez-ortega *et al*. 2011; Wu *et al*. 2018). Most of HIV variation in untreated patients has been argued to arise from mutations gradually compensating the early escape mutations conferring resistance to the immune response (Rouzine and Coffin 1999).

A large number of approaches have been proposed to measure epistasis from genomic data (Cordell 2009; Chen *et al*. 2011; Ueki and Cordell 2012). The simplest methods are based on pairwise allelic correlations (Hill and Robertson 1966; Barton 1995). The problem with all these approaches is that linkage and indirect interactions, create strong inter-site correlations even between non-interacting locus pairs, and these false-positive pairs are much more numerous than the true epistatic pairs. Stochastic effects are well-recognized as the most serious obstacle to the detection of epistatic effects by any method (Wei *et al*. 2014). In a single asexual population, stochastic linkage overshadows the epistatic footprint, except in a narrow range of times and parameters (Pedruzzi and Rouzine 2019). The same limitation exists for tree-based methods of epistasis measurements (Kryazhimskiy *et al*. 2011; Neverov *et al*. 2015).

A method to filter out false links due to indirect interactions has been developed and applied to protein sequence sets obtained from different species (Weigt *et al*. 2009; Cocco *et al*. 2018). The technique is based on the combination of a covariance method with the estimates of the direct mutual information. The true interactions are inferred from the condition that entropy (defined as the probability of configuration or the degree of disorder) is maximal given the observed frequencies of single sites and site pairs. A similar technique has been used to measure the fitness landscape of Ab binding regions of HIV surface protein gp120 (Louie *et al*. 2018). While this technique eliminates indirect interactions in fairly complex network, it is not designed to exclude stochastic linkage effects. The present work is designed to fill this gap.

Recently, we proposed an evolution-based approach to the problem of detection of epistatic effects (Pedruzzi *et al*. 2018), which we describe here, since it is relevant for our method detailed in *Results*. We simulated the evolution of a pair of loci within a multi-locus genome in an adapting population. We demonstrated the existence of a broad parameter and time range, where each fitness class has sufficient time to assume the most probable state, which changes slowly in time, as the traveling wave shifts in time. This situation will be referred below as “quasi-equilibrium”. The time required for the system to arrive there is *t* ≫ 1/ *⟨s⟩* (Pedruzzi *et al*. 2018). In quasi-equilibrium, the entropy *S*(*t*) (degree of disorder) is at its current maximum and represents a function of fitness *W*(*t*), which slowly changes in time during the process of adaptation: *S*(*t*) = *F*[*W*(*t*)]. The following derivation employs the fact, predicted by all the traveling-wave models cited above, that the distribution of fitness *W* is narrow, as given by std[*W*] ≪ *W*. Quasi-equilibrium is possible due to a very slow motion of the wave, limited by the generation and establishment of new beneficial mutations at the front edge of the wave (Rouzine *et al*. 2003; Desai and Fisher 2007; Neher *et al*. 2010; Hallatschek 2011). We consider the regime of adaptation, where the population is very far from full equilibrium (steady state), so that we can neglect with deleterious mutation.

Consider a pair of loci (Fig. 1A) and assume that each can have one of two variants, alleles, either beneficial or deleterious allele, denoted 0 or 1, respectively. We denote with *s*_1_, *s*_2_ the respective fitness effects of mutations. There are four possible haplotypes, 00, 01, 10, 11. We assume that the two sites interact epistatically. If both deleterious alleles are present, they partly compensate each other to the degree *E*, where 0 < *E* < 1. Then, the log fitness values of the four haplotypes are given by

**Fig. 1.**
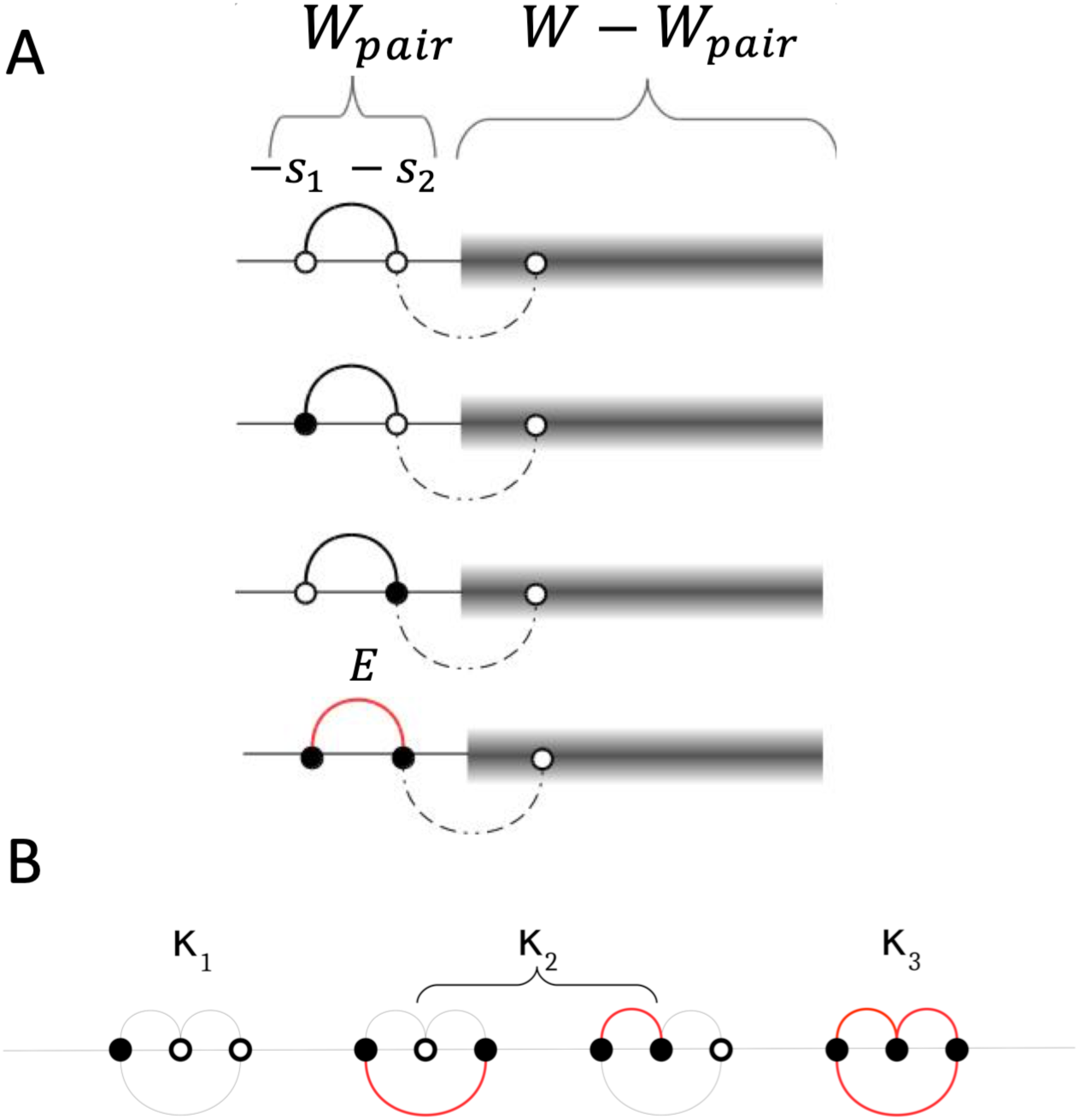
(A) A pair of interacting sites in a long genome. Black and white circles are deleterious and favorable alleles, respectively, arches show direct (potential) epistatic interactions, red arches are actual interactions between deleterious alleles. Dashed line: neglected interaction. Grey box: the rest of the genome. (B) A simplest epistatic network illustrating the effect of indirect interactions.. Numbers *k*_*i*_ denote the numbers of clusters with *i* connected sites in a genome.

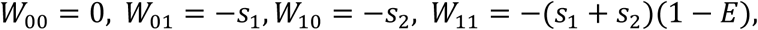

respectively. If *E* is allowed to change below 0 or above 1, we can obtain any type and sign of epistasis: positive, negative, and overcompensation.

The frequency of each haplotype *ij* in a population, where *i* and *j* are 0 or 1, is denoted *f*_*i j*_. As is easy to understand, *f*_*i j*_ is proportional to the number of sequence configurations of the rest of the genome, denoted exp(*S*_*rest*_) (grey rectangles in Fig. 1A). Entropy *S*_*rest*_ depends on the fitness of the rest of genome equal to *W* − *W*_*i j*_. Hence, the entropy of each haplotype is *S*_*rest*_ = *S*(*W* – *W*_*i j*_). Since the genome is long, *W*_*i j*_ is much smaller than *W*, so we can expand linearly

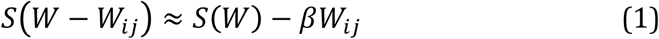

where *β* = -*dS*/*dW* > 0. Thus, the frequency of each haplotype *f*_*i j*_ is proportional to the corresponding configuration number, exp *S W* − *W*_*i j*_. Hence, we obtain

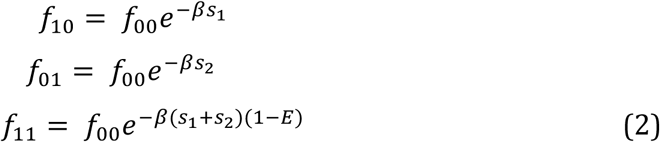

where *β* depends on *W* and, hence, on time, but is the same for all loci (Pedruzzi *et al*. 2018). This simple derivation neglects interaction of the site pair with the other sites of the genome. This assumption can be lifted and generalized for any interaction network (Pedruzzi *et al*. 2018). We also do not need to restrict ourselves to the bi-allelic model, but we will keep this assumption, since it is works in the short term. The Boltzman factor in Eq. 2 was derived originally for a small subsystem connecting to a large thermal reservoir. We remind, however, that we are not discussing the state of equilibrium, but adaptation far from equilibrium.

Eq. 2 predicts, in the general form, how the average haplotype frequencies evolve in time. It is tempting to use it for estimating parameters *E, s*_1_, *s*_1_ from the observed haplotype frequencies, and thus find the fitness landscape for the entire genome (at least, if epistasis is pairwise, as we assumed). In reality, these formulae are not readily applicable to real-life sequences. because of the strong random variation of *f*_*i j*_ caused by stochastic linkage effects we have discussed. Of course, the problem of strong linkage noise causing poor reproducibility of various methods [see (Wei *et al*. 2014) for review] is not specific for this method of estimating epistasis. Any attempt to measure epistasis in a single population, whether by measuring covariance from measure *D*′, *r*^2^, mutual entropy, UFE (Pedruzzi *et al*. 2018) or by tree-based methods (Neverov *et al*. 2015) faces the same problem.

The epistatic pairs in a single population are not observable in principle, due to the phylogenetic relationship of sequences in population. The problem is that all the sequences in a population closely resemble the common ancestor, which diverges from the origin in a random direction (Pedruzzi and rouzine 2019). As a result, any measure of co-variance, or even the use of the entire tree, produces only strong noise of random sign. Co-variation due to random linkage completely masks the epistasis signature in a population. The only way to resolve this issue is by averaging over many independent populations with similar parameters under similar conditions.

Thus, without sampling multiple populations, it is not possible to infer epistasis in principle, due to the stochastic nature of phylogenetic relation of sequences. This fundamental limitation cannot be resolved by any existing or future method. The contribution of the present work is not to overcome this limitation, which is not possible in principle, but to demonstrate the existence of a large residual stochastic errors left after averaging over a limited ensemble of populations, and to propose a new method to compensate this error. Here we offer a new technique, based on the application of the quasi-equilibrium argument (above) to triple-way haplotypes. We show analytically and by simulation that a triple-way haplotype test represents an effective way to eliminate residual false links, including linkage and indirect interactions. We will demonstrate its high fidelity in a broad parameter range on the mock sequences obtained by computer simulation.

After training the method on simulated sequences, we apply this technique to real virus sequences from an adapting population. Influenza virus evolving in a population was demonstrated to obey the traveling wave theory with an effective selection pressure caused by the accumulating memory cells (Rouzine and rozhnova 2018; Yan *et al*. 2019). Therefore, it is expected to be amenable to our method. Using 8000 influenza sequences obtained from various geographic locations sequences, both before and after the pandemic of 2009, we infer the epistatic network in a surface protein, neurominidase, by comparison of the old and new strains. We chose this specific protein, because it affects infectivity and underwent strong changes when it was replaced by a new strain in 2009. It also has been a target for drug therapy.

We note that the old strain and the pandemic strain of H1N1 share 80% of homology. This indicates that, at some point in the past, the two strains had a common ancestor, and later they diverged due to accumulation of mutations. Our aim is find out now how the new sequence variant can obtained by changing the old variant by making multiple mutations, including primary and secondary mutations, without much loss of fitness. Our method does not depend on the actual path of the divergence of the two variants, nor can it infer the order in which these secondary mutations have emerged, but it can infer the resulting network. The same general approach is routinely used for estimating the epistatic networks from the comparison of proteins of strongly divergent species with presumably weak linkage effects (Weigt *et al*. 2009; Cocco *et al*. 2018). The difference of our approach is that it filters out false-positive links due to linkage, as well as indirect interactions, rather than indirect interactions alone.

## RESULTS

### Simulation model to generate sequences for the test

We start by simulating the stochastic asexual evolution of a haploid population using a Wright-Fisher process in the presence of the factors of random mutation, random genetic drift, and directed constant selection (Fig. 2b) (*Methods*). We assume two possible alleles at each locus (site), 0 and 1. The binary simplification provides a major reduction in the computational cost and is especially accurate for relatively conserved sequences. We set some pairs of sites to interact epistatically, with a mutual degree of compensation between deleterious alleles chosen to be *E* = 0.75. It represents an example of strong compensation, 75%. At lesser *E*, the detection of epistatic links is worse. We consider a simple epistatic network consisting of double arches (Fig. 1B). The choice of the network topology does not affect the detection power of our method. Indeed, as we show below, most false-positive interactions in the short-term evolution are due to stochastic linkage effects rather than indirect epistatic interactions.

**Fig. 2.**
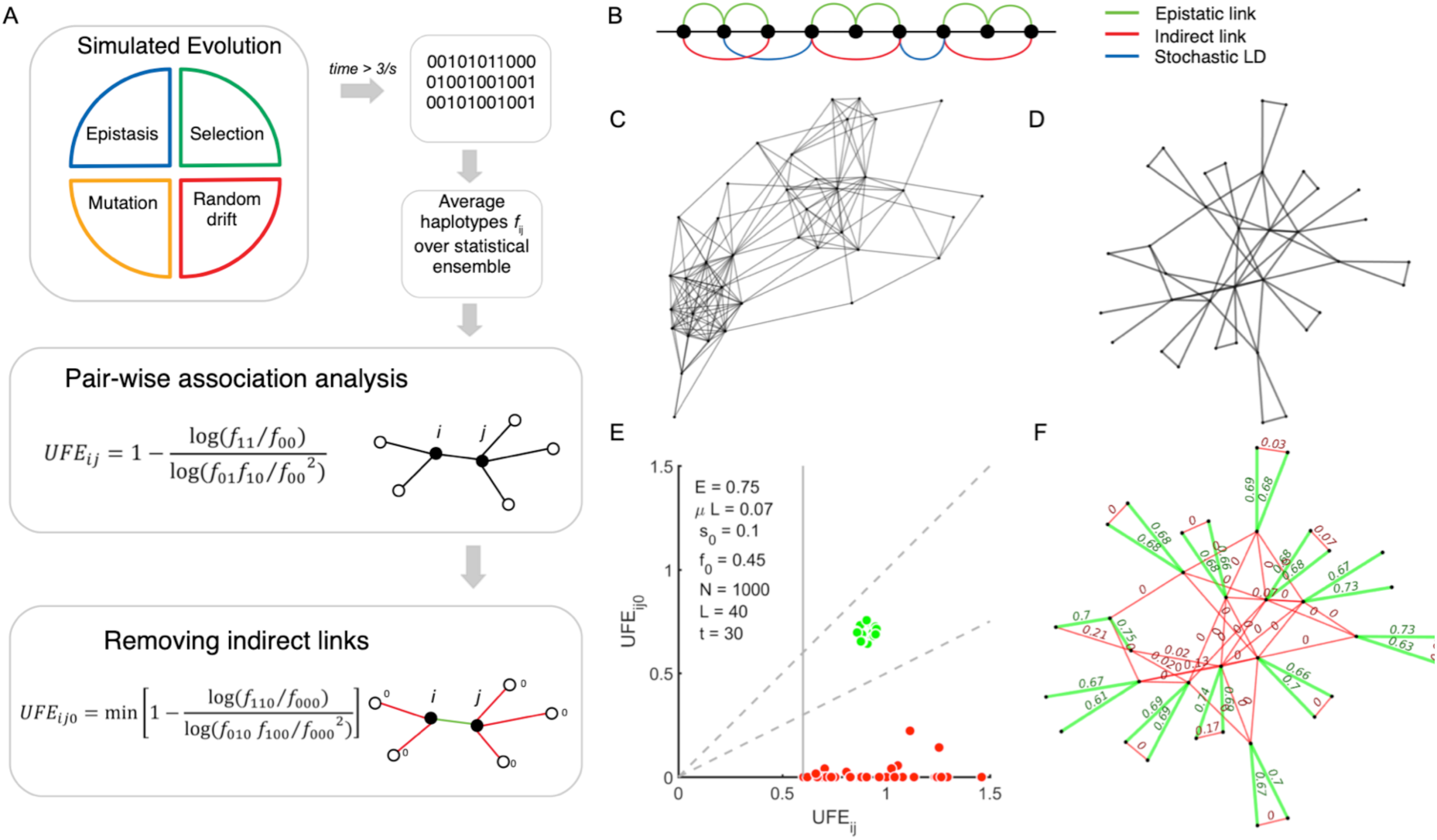
Schematic diagram of the method and its training on simulated sequence data. **A**. The computer model of asexual evolution includes the factors of random mutation, selection, epistasis, and random genetic drift. Pairwise haplotype frequencies *f*_*ij*_ are averaged over an ensemble of independent simulation runs (populations). The strength of interactions UFE_*ij*_ is calculated. The indirect links and the residual stochastic LD are excluded by using triple-site haplotype frequencies, UFE_*ij0*_. **B**. Pre-set epistatic network for 50 sites. Real epistatic links are shown by green lines. The resulting indirect links are red lines. Some examples of stochastic linkage bonds are shown by blue lines. **C-D**. The network of strong (UFE > 0.5) candidate epistatic interactions predicted (**C**) from a single population and (**D**) after averaging over 200 populations. **E**. Scatter plot of 3-locus haplotype min(UFE_*ij*0_) shown against UFE_*ij*_ for the pairs identified in (**D**). Dashed sector (green): Predicted direct interactions. **F**. Predicted network accurately recapitulates the pre-set epistatic network. Parameters: initial allele frequency *f*_0_ = 0.45, mutation rate per genome *μL* = 0.07, fixed selection coefficient *s* = 0.1, *N* = 1000, *L* = 40, epistatic strength *E* = 0.75.

### First step: averaging over populations

Genome sequences produced by the simulation demonstrate the presence of strong linkage disequlibrium (LD) originating from three sources: direct epistatic interaction, indirect interaction, and stochastic linkage effect. Out first task is to try to detect possible epistatic interactions from the simulated sequences using pair-wise association analysis (Fig. 2A). As we mentioned in Introduction, their detection is masked by strong stochastic linkage arising from phylogenetic divergence of independent populations (Pedruzzi and Rouzine 2019). To decrease the linkage effect, we calculate pairwise haplotype frequencies, denoted *f*_ij_, for each pair of sites and average them over multiple evolutionary-independent populations of the same size. Then, we perform a pairwise association analysis using a correlation measure we termed “Universal Footprint of Epistasis” (UFE), which is a direct estimator of the degree of mutual compensation of two deleterious alleles, *E* (Pedruzzi *et al*. 2018) (*Methods*). More traditional LD measures, including *D’* and Pearson coefficient *r*^2^, have been shown to generate similar stochastic error (Pedruzzi and Rouzine 2019). We select pairs with high correlation, UFE > 0.6. Before we average over populations, for a single population, the raw cluster of inferred pairs is extremely complex and completely hides true epistatic interactions (Fig 2C). A significant reduction of the number of false-positive bonds is obtained by averaging *f*_ij_ over 200 independent populations (Fig. 2D). However, the fast majority of remaining links are still false-positive, since their number is much higher than the number of actual interactions (Fig. 2B, green arches).

### Second step: Triple-site haplotype method

Although much simpler, the resulting network in Fig. 1D is still densely populated by residual linkage links and indirect links, which are as strong as direct interactions. To detect the residual false links, we use a new procedure based on the quasi-equilibrium approximation. We test the validity analytically in *Methods*, as well as below by the direct comparison of the inferred network with the network set in simulation. The procedure is, as follows.

For every potential link between sites *i* and *j* detected in Fig. 2D, we recalculate the haplotype frequencies and UFE, by including only the sequences that contain 0 at one of the adjacent connected nodes of the link and denote it UFE_*ij*0_ (Fig. 2A, bottom). In other words, we calculate the frequencies of three-way haplotypes 110, 100, 010, and 000. Considering different 0-nodes for all connected nodes of the link, we then find the minimum value of UFE_*ij*0_ over all these nearest-neighbor nodes. The scatter plot in Fig. 2E demonstrates that, for the false pairs, min(UFE_*ij*0_) is several-fold smaller than UFE_*ij*_ (red dots in Fig. 2E). For true links (green dots in Fig. 2B,E), the two correlation measures are nearly the same. The reason for this difference is that a 0-node removes a detour path around the link that causes indirect correlations, thus decreasing the false association of the alleles. An analytic demonstration is given in *Methods*. Therefore, we identify and remove false links as those with low min(UFE_*ij*0_)/UFE. The choice of the threshold is not crucial, as long as we average *f*_*ij*_ over, at least, 20 populations for our model parameter choice, so that the two groups, false and true interactions, remain distinct. As a result, for our example, we obtain 100% perfect detection (Fig. 2B). As a bonus, we obtain accurate estimates for the compensation strength, *E*, because UFE ≈ *E* within 15% accuracy (Fig. 2F, numbers at green links). We can conclude that the triple-haplotype method, for the parameters tested, has a high fidelity with simulated sequences with a known epistatic network.

### Application to influenza A virus

After training our method on simulated sequences, we calculate the interaction network for real virus sequences isolated from an evolving population. Our choice is the surface protein of Influenza A virus strain H1N1, Neuraminidase (NA), known to be an important target of drug therapy and immune response. This protein is one of two proteins that control the virus entry into a host cell, the other being Hemaglutinin. Our aim is to identify epistatic network responsible for the difference between the pre-2009 strain and the pandemic strain of 2009. For this aim, we compare the sequences of the first strain to the sequences of the second strain sampled worldwide.

We downloaded 8440 sequences of NA from a public database (https://www.fludb.org). They were collected worldwide, from different geographic locations, between years 2005 and 2010, and included both pre-pandemic and post-pandemic strain. All sequences were aligned. Then, we found the consensus (majority) allele for each aminoacid position. To simplify our task, we binarized the sequences by setting each consensus allele to 0 and each non-consensus (“mutant”) allele to 1. This simplification is adequate for the aim of detection of interactions, unless several aminoacid variants are present at a site at similar frequencies, which we found to be a rare occurrence.

We observed a bimodal distribution of sequences in the mutant allele frequency per genome with two separate maxima of different height. The low-frequency peak was taller. The bimodal distribution reflects the mixture of two strains, the old and the new, with 80% homology. In order to compensate for unequal sampling from the early strain and the late strain, the more abundant sequences with mutation frequency per genome less than a preset value, *d*_v_, were randomly sampled and down-weighted by a coefficient, *D*_*w*_, ranging from 5% to 50%. To obtain the average pairwise haplotype frequencies, *f*_*ij*_, we repeated the resampling multiple times. Then, we followed the procedure described above (Fig. 2C-F) to infer the intra-protein network of interactions (Fig. 3).

**Fig. 3.**
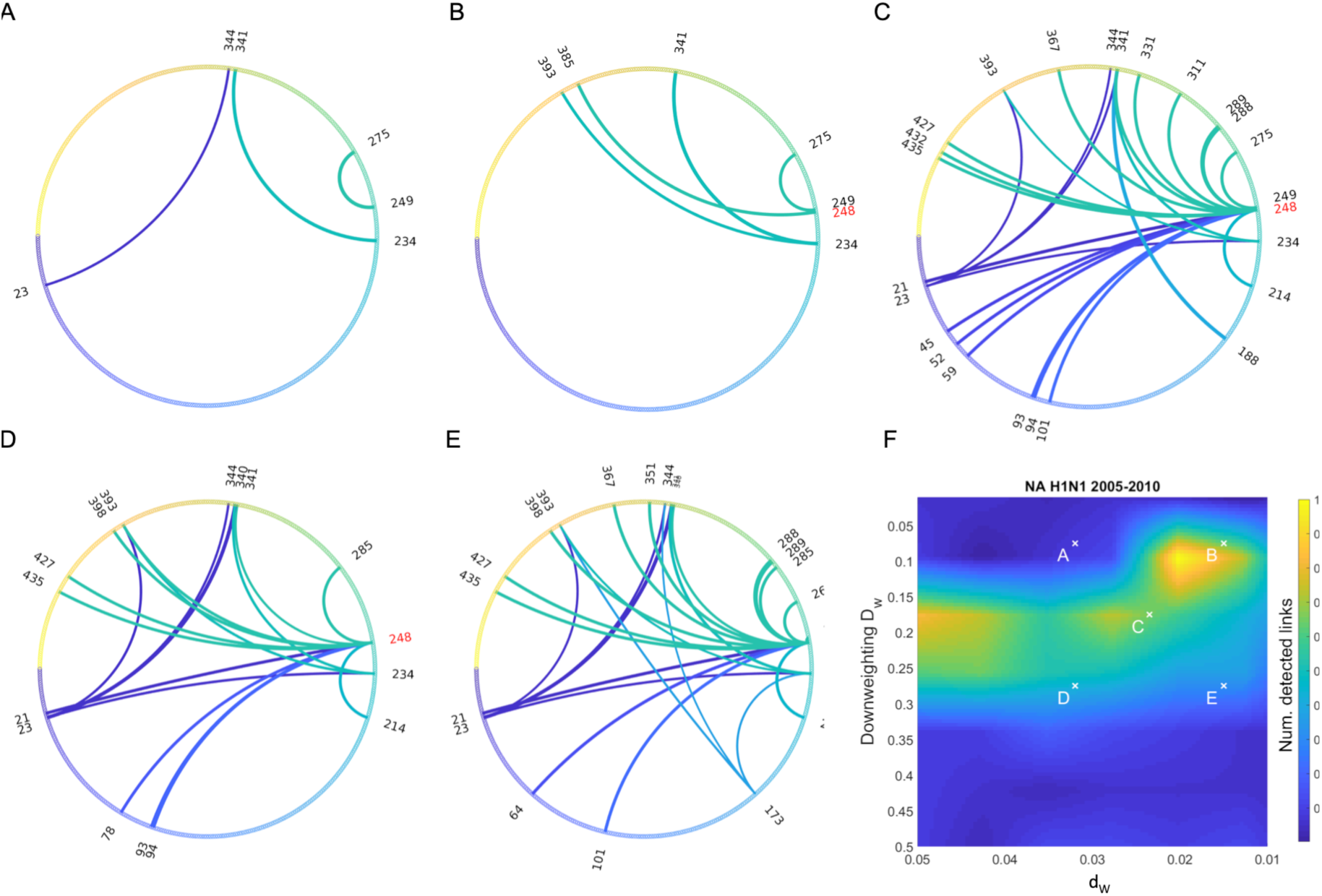
Epistatic network predicted from sequence data on surface protein sequences of Influenza A H1N1 obtained in 2005 and 2010. The circular diagrams show the network of interaction for neuraminidase. Sequences with homology to the consensus less than *d*_*v*_ are randomly re-sampled many times, with their number downweighted by coefficient *D*_*w*_. (**F**) 2D heatmap showing the total number of links as a function of *d*_*v*_ (X-axis) and *D*_*w*_ (Y-axis). Different versions of wheels in **A-E** correspond to different choices of *D*_*w*_ and *d*_*v*_ shown by crosses in **F**.

### Sensitivity to sampling parameters

The resulting network is moderately robust to the variation of (*d*_*v*_, *D*_*w*_) (Fig. 3A-F). There exists a plateau region of (*d*_*v*_, *D*_*w*_) where the total number of links varies weakly (Fig. 3F). We observe that site 248 in NA represents the primary site connected to multiple compensatory sites (Fig. 3D). The dependence of results on (*d*_*v*_, *D*_*w*_), which infers between 15 and 22 compensatory sites, probably originates from the unequal presentation of local subpopulations in the database. As a result, stochastic linkage effects average out less than optimally. Below, we choose the network variant shown in Fig 3D as the “golden middle” of the set.

### Structural interpretation

The inferred epistatic sites are shown on the three-dimensional protein structure of NA. The active pocket of NA (purple) serves to bind sialic acid on target cell surface (Fig 4A). This result shows that the inferred primary mutation at residue 248 is located near the active pocket. The inferred compensatory mutations (Fig. 3D) helping NA to restore and improve its fitness are located on protein surface inside alpha-helixes, determining the mutual orientation of beta-sheets. Experimentally, mutation 248 is well-known. It was shown to enhance the low-pH stability of NA (Takahashi *et al*. 2013) and is found in all influenza A H1N1 strains after 2009 pandemic, regardless of a geographic location (Seibert *et al*. 2012; Byarugaba *et al*. 2016; Otte *et al*. 2016).

**Fig. 4.**
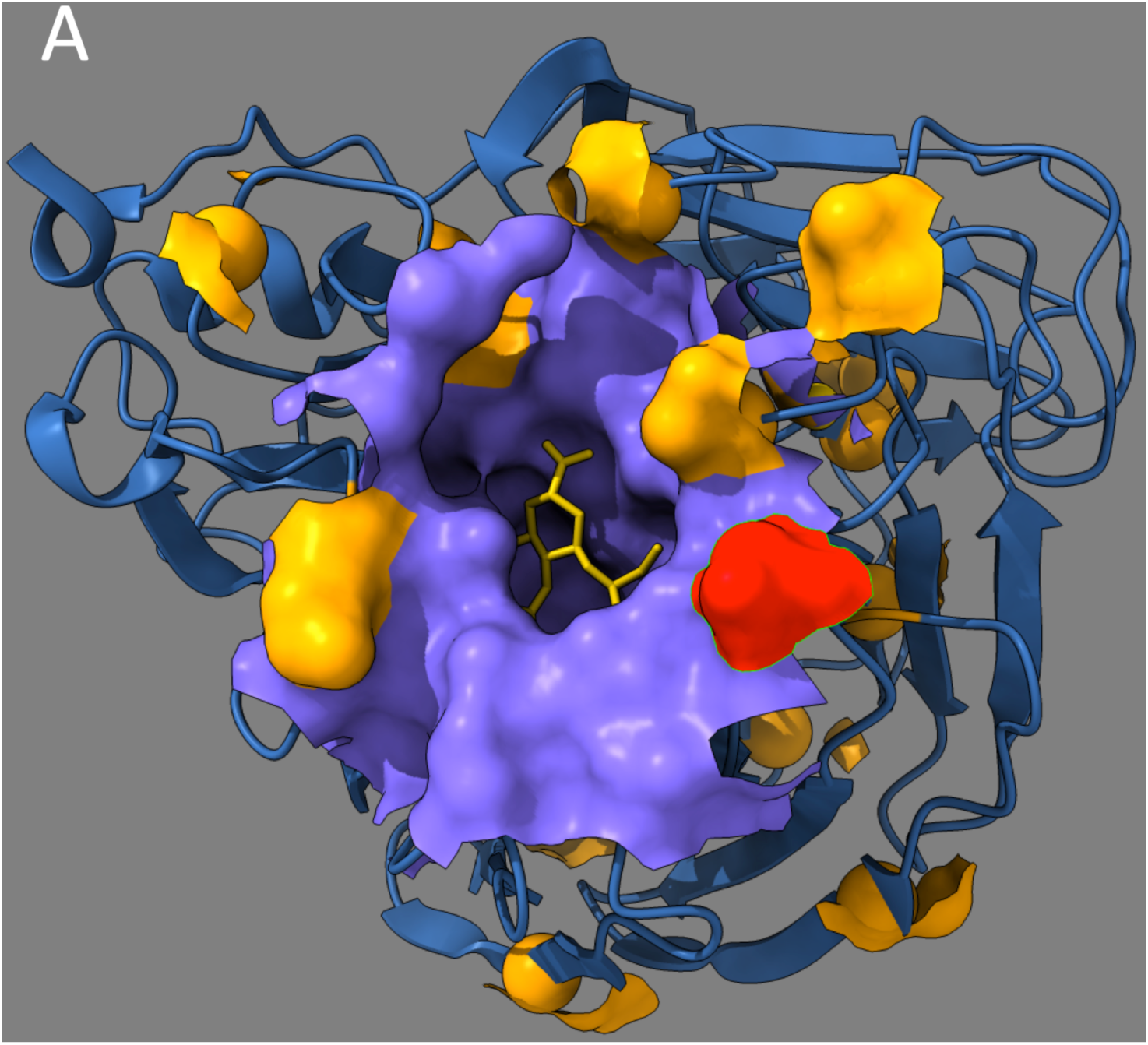
Structural location of the predicted epistasis network for the neuraminidase of influenza virus. The figure shows the three-dimensional structure of Influenza A H1N1 neuraminidase (PDB ID code 4QVZ). Colored spheres represent predicted epistatic residues from Fig. 3D. Red sphere: Predicted primary mutation (residue 248 in Fig 3D). Orange spheres: Compensatory residues from Fig. 3D.

## DISCUSSION

In the present work, we propose an efficient evolution-based method to identify the co-variance caused by epistasis from the co-variance caused by stochastic linkage effects and indirect interactions. First, we average the observed haplotype frequencies over independent populations, then we select the links with a high co-variance, and then we apply a tri-way haplotype test for each candidate bond to eliminate the effects of linked sites due to common inheritance.

The tri-way haplotype method is justified using a simple analytical model (*Methods*) assuming a quasi-equilibrium state created by a slowly-moving fitness wave (not to be confused with quasi-linkage equilibrium or mutation-selection balance). The existence of quasi-equilibrium has been tested previously by simulation in a broad parameter range (Pedruzzi *et al*. 2018; Barlukova *et al*. 2020). Intuitively, because population evolves slowly, the distribution of alleles between sites has sufficient time to obtain the most probably, most disordered state given a current fitness.

We use a mock sequence set evolved in a Wright-Fisher population with a known epistatic network to demonstrate the high fidelity of the method in a controlled environment. The method eliminates false bonds caused by both indirect interactions and linkage, at least, in the case of a relatively simple epistatic network and moderately diverse populations (< 40% diversity).

We applied this technique to identify primary and secondary mutations in influenza A H1N1 neurominidase, responsible for the difference between the pre-2009 and post-2009 variants. We do not address the origins or the history of the strain. Influenza virus has been shown to map to the traveling wave theory (Rouzine and Rozhnova 2018; Yan *et al*. 2019), which justifies the use of the quasi-equilibrium assumption. Our method infers a single primary site and 15-20 strong multiple compensatory mutations, which number is in the same general range as the number of compensatory mutations observed, for example, in drug-resistant strains of HIV. The inferred primary mutation, 248, has been observed in all influenza A H1N1 strains after 2009 pandemic in various geographic locations (Seibert *et al*. 2012; Byarugaba *et al*. 2016; Otte *et al*. 2016). I was shown to affect virus infectivity (Takahashi *et al*. 2013). Therefore, our method is capable of pinpointing the well-known primary mutation responsible for the emergence of the pandemic strain.

As compared to the existing technique of elimination of indirect interactions developed for different animal species, which diverged for millions of years (Weigt et al. 2009; Cocco et al. 2018), our method is designed to be used for recently diverged populations (thousands of generations or less) of the same species with small to moderate diversity. We have tested it for Hamming distance *2f*(1 − *f*) < 0.5. Furthermore, our method is capable of eliminating stochastic linkage, less important when comparing different species (Weigt et al. 2009; Cocco et al. 2018). Where both methods can be potentially applied, such as calculating the fitness landscape of HIV Ab-binding regions (Louie *et al*. 2018), our method is much faster computationally, because it is local in the genome. Indeed, we can consider one pair of loci at a time and not to worry about simultaneous optimization of *L*^2^/2 parameters of the full interaction matrix. Also, it helps to avoid the situation when the number of fitting parameter is too large, and the system is over-defined.

To illustrate application our method, we averaged haplotype frequencies over influenza H1N1 sequences obtained from a large number of geographic locations. Influenza sequences represent a meta-population with many islands, not a single well-mixed population. The assumption here is that that the averaging over different geographic location allows to obtain, at least, a partial ensemble average. It is unclear to which extent the lack of the full ensemble average affects the estimate, but at least, we are able to compensate the residual linkage errors within that partial ensemble, something that has not been done before.

The main limitation of the proposed approach is that it assumes constant directional selection, as opposed to balancing selection or time-dependent selection, such as occurs under changing external conditions or, in the case of pathogens, under the rising immune response. While the specific case of influenza evolving in a population under accumulating immune memory B cells has been shown to map to the case with constant selection (Rouzine and Rozhnova 2018; Yan *et al*. 2019), the case of virus evolution within an animal under CTL response is less trivial (Batorsky *et al*. 2014). Both cases require further investigation.

To summarize, we proposed a technique to infer the epistatic effect on evolution of locus pairs and tease it out from stochastic linkage effects. We hope that our quasi-equilibrium approach and further development of this technique will prove useful for all researchers interested in finding fitness landscapes of various organisms from their genetic samples. Also, it will be very valuable to compare these data with the result of deep mutational scanning (Lee *et al*. 2018; Hom *et al*. 2019), which we hope to do in the future.

## METHODS

### Wright-Fisher model for simulation

We simulate the evolution of a haploid asexual population of *N* binary sequences. In an individual genome, each locus (site, nucleotide position, aminoacid position) numbered *i* = 1, *2*, …, *L* is occupied by one of two alleles, either the wild-type allele, denoted *K*_*i*_ = 0, or the mutant allele, *K*_*i*_ = 1. We use a discrete generation scheme in the absence of generation overlap (Wright-Fisher model). The evolutionary factors included in the model are random mutation with rate *μL* per genome, constant directed selection, and random genetic drift due to random sampling of progeny. Selection includes an epistatic network with a set strength and topology. Recombination is absent. A previous modeling study shows that moderate levels of recombination can enhance epistatic detection (Pedruzzi and Rouzine 2019). Where use the standard model of fitness landscape with pairwise interaction. The logarithm of the average progeny number, *W*, depends on sequence [*K*_*i*_] as given by

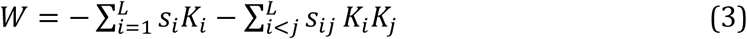

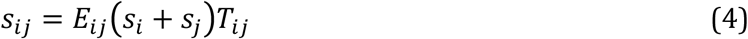

Here the selection coefficients *s*_*i*_ and *s*_*i*_ denote the fitness costs of two deleterious mutations that are partially compensated by each other. By definition, *E*_*ij*_ is the degree of compensation of deleterious alleles at sites *i* and *j*. Values *E* = 0 and 1 represents no epistasis and full compensation, respectively.

In our example in Fig. 2, we consider a haploid population with initial frequency *f*_0_ = 0.45, mutation rate *μL* = 0.07, fixed selection coefficient *s* = 0.1, *N* = 1000 individuals, and *L* = 40 sites, and fixed epistatic strength *E* = 0.75.

### Quasi-equilibrium approximation

In a broad parameter range, an adapting population represents a slowly-moving, narrow peak in fitness space (Rouzine *et al*. 2003; Rouzine and Coffin 2005; Rouzine *et al*. 2008; Rouzine and Coffin 2010; Good *et al*. 2012). Evolution is slow, because the limiting factor is the addition of a rare beneficial mutation established within a highly-fit genetic background (Rouzine *et al*. 2003; Good *et al*. 2012). Because the fitness distribution moves slowly, the entropy (disorder degree) of the mutation distribution over genomes has enough time to reach its current maximum, restricted by the current average fitness of the population. This situation is called “quasi-equilibrium”. At each moment, each fitness class has enough time to reach the most probable, most chaotic state given its fitness. Previously, we verified the validity of quasi-equilibrium in a broad range of parameters and initial conditions after time ∼ 1/<*s*> (Pedruzzi *et al*. 2018). When the population finally arrives in mutation-selection equilibrium, the presence of deleterious mutation events compensates the natural selection and creates linkage equilibrium. At this point, the method breaks down.

### Linkage measure

Eqs. 1 and 2 can be used to estimate epistatic interaction. We will use a binary measure of allelic correlations (Pedruzzi *et al*. 2018)

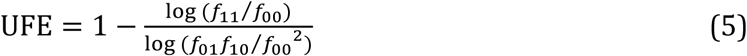

where *f*_00_, *f*_10_, *f*_01_*f*_11_ are the haplotype frequencies averaged over ensemble.

UFE performs similarly to more traditionally used measures, such as Lewontin’s *D’* and Pearson correlation coefficient, *r*^2^ (Pedruzzi and Rouzine 2019) or mutual information (Weigt *et al*. 2009; Cocco *et al*. 2018). As compared to these measures, UFE has the unique advantage of directly measuring the degree of mutual compensation of two alleles *E*, provided they are epistatically isolated. If the locus pair does not interact with the other sites in the genome, we have UFE = *E* (see Eqs. 1 and 2). If they are a part of a network, this measure overestimates *E*. In the main text, we calculate UFE for every pairs of sites (Fig. 2A). We leave only those pairs where UFE exceeds a set threshold of 0.6.

### Tri-site test of false bonds

To test whether a detected correlation for a pair of sites is direct, and not due to linkage or indirect correlation, for each connected pair *i, j*, we also calculate the three-way measure

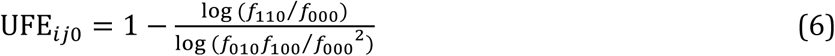

where 0 in the third position selects only for the sequences with the consensus allele 0 at a chosen site connected to one site of the tested pair. We consider all possible connected sites as 0-nodes and calculate the minimum value of UFE_*ij*0_ over all possible 0-nodes. This method cuts the most important indirect correlation by a “detour” for a site pair. It works for both linkage and indirect correlation.

### Test of the method using a simple analytic approach

To illustrate how the above method works on indirect bonds, let us consider a simple analytic model from (Pedruzzi *et al*. 2018). We assume that some sites carry deleterious alleles with equal selection coefficient *s*_1_ = *s*_2_ and a fixed epistatic strength *E*_*i j*_ = *E* (Eq. 3) In this simple case, we can fully characterize a genome by the numbers of interacting allelic clusters of different size. Let *k*_*i*_ denote the number of clusters with *i* nodes and *b*_*i*_ bonds. Then, from Eq. 3, we can express log fitness *W*as a sum over clusters of different size (Fig. 1B)

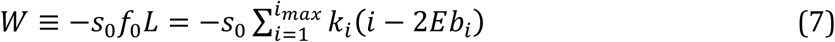

New notation *f*_0_ has the meaning of the effective frequency of uncompensated mutations that would have the same total fitness *W*. The number of bonds *b*_*i*_ At *i* ≥ 3 depends on the topology of epistatic network, while *b*_1_ = 0, *b*_2_ = 1 always. In our example with the network of double arches, we have *b*_3_ = *2* (Fig. 1B).

As we mentioned above in *Intro* and *Methods*, we assumed the state of quasi-equilibrium (not to be confused with steady state) determined by the current fitness. Therefore, at each moment of time, numbers *k*_*i*_ are determined by the condition that the entropy of the system is maximum given fitness, Eq. 7. Entropy *S* is defined as the log number of sequence configurations

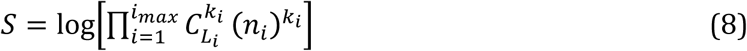

where *L*_*i*_ is the number of all possible locations for a cluster of size *i*, and *n*_*i*_ is the number of each cluster’s configurations (shapes). The values of *L*_*i*_ and *n*_*i*_ depend on the network topology.

Previously, we applied this argument for several topologies to derive the numbers of clusters of different size, as follows (Pedruzzi *et al*. 2018). We showed, for the one in Fig. 1B, that the numbers of clusters of size *i* = 1, *2* and 3 in, normalized as *f*_*i*_ = *k*_*i*_/*L*, are related as

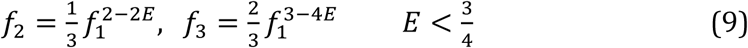

The 1st and 3d site in each triplet in Fig. 1B do not interact directly, but only indirectly through site 2. For these two sites, the haplotype frequencies are

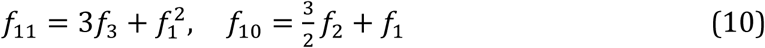

When epistatic interaction is sufficiently strong, as given by the condition *E* > 1/2, triplets dominate over single alleles and their doubles, as given by *f*_1_ ≪ *f*_2_ ≪ *f*_3_ (Eq. 9). In this case, we can approximate haplotype frequencies as

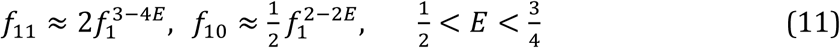

Using covariance measure UFE_*i j*_ = defined in Eq. 3, we obtain 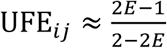. We observe that the indirect covariance between sites 1 and 3 is as strong, ∼ 1, as for directly interacting sites. For directly interacting sites 1 and 2 (Fig. 1), we previously obtained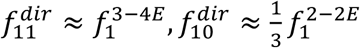, see (Pedruzzi *et al*. 2018),

*Supplement*, Eqs. (3.29) and (3.30).

However, if we calculate UFE _*i j 0*_ instead of UFE _*i j*_ by including only the sequences with majority allele 0 at site 2, we obtain

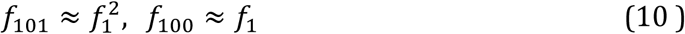

instead of Eq. 9, and Eq. 6 yields UFE _*i j 0*_ = 0. Thus, the phantom covariance disappears, when we select only sequences with a majority allele inserted between the two sites tested. This result is intuitively clear: by the definition of fitness (Eq. 1), only minority alleles interact with each other, while majority alleles form neutral background. The same method turns out to be extremely effective for eliminating false bonds created by linkage (compare Fig. 2B vs 2E).

### Sequence preparation

We have applied the three-way test to influenza virus. We performed a multiple progressive alignment for amino acid sequences obtained for a surface protein of Influenza A virus strain H1N1, Neuraminidase (NA), from public database https://www.fludb.org. We focused on NA because of the massive amount of sequence data, because it was a target of drug treatment, and because strong changes in NA were responsible for the higher infectivity of pandemia in 2009.

We have used about 8000 sequences found in the cited database for the period 2000-2010 from various geographic locations. Our aim was to understand the differenc between the protein variants before and after pandemic of 2009, which have 80% of mutual homology. We aimed at discovering only the epistatic sub-network related to that difference, and were not interested in any other epistatic interactions in these proteins. For this end, we compared worldwide samples of sequences from the two strains. We randomly sampled similar amounts of sequences from the first and second strains, and re-sampled them several hundred times. We also checked the robustness of the results to exact sampling size (Fig. 2F, *Results*).

Pairwise distances between sequences were computed using pairwise alignment with the Gonnet scoring matrix implemented in MATLAB. To calculate the guide tree we used the neighbor-joining method assuming equal variance and independence of evolutionary distance estimates. The obtained consensus served as a universal reference to binarized data sequences. Before applying the detection algorithm, the protein sequences were binarized, by direct comparison of each sequence to the consensus. Each amino-acid residue was set to 0 or 1 for consensus or non–consensus. Although combining all aminoacid variants per site ignores the specific biochemistry of substitutions, this approach greatly reduces the number of haplotype combinations and also increases the sensitivity by effectively increasing the haplotype frequencies.

Next, we measured the mutational frequency for each sequence along sequences and for each site across sequences. The subset of low-diversity sequences with allelic frequency below a cut-off *d*_*v*_ was randomly sampled and down-weighted according to a set coefficientm, *D*_*w*_. Then, we determined the average pairwise and three-way haplotype frequencies for all pairs and triplets of sites, as described above.

### 3D structures

To characterize the three-dimensional network of epistatic interaction in Fig. 4, we used software package ChimeraX from internet site https://www.rbvi.ucsf.edu/chimerax/.

## Data availability

Influenza sequence data are from public database https://www.fludb.org.

## Acknowledgements

We thank Martin Weigt and Alessandra Carbone for useful comments. This research has been funded by Agence Nationale de la Recherche grant J16R389 to IMR, http://www.agence-nationale-recherche.fr/.

